# Imipramine binds to amyloid-β and extends the survival of Alzheimer’s disease patients

**DOI:** 10.64898/2026.10.28.762584

**Authors:** Noah R. Johnson, James Beham, Stefan Sillau, Juliana Read, Athena Ching-Jung Wang, Md. Mahiuddin Ahmed, Daphne Quang, Daniel Stephenson, Morkos A. Henen, Beat Vögeli, Heidi J. Chial, Robb Welty, Liliya Vugmeyster, Huntington Potter

## Abstract

**Background:** Apolipoprotein E (apoE) plays numerous roles in the pathogenic pathway to Alzheimer’s disease (AD). We previously performed a high-throughput screen and identified the antidepressant imipramine as an inhibitor of the interaction between apoE and amyloid-β (Aβ). We also found that imipramine use to treat depression in AD patients was associated with better Mini-Mental State Exam (MMSE) scores. Here we studied the mechanisms by which imipramine inhibits the AD pathogenic pathway and investigated its effect on mortality in AD patients.

**Methods:** Nuclear magnetic resonance spectroscopy was used to characterize the interaction between imipramine and Aβ monomers. Human induced pluripotent stem cell (hiPSC)-derived AD cerebral organoids (COs) and transgenic AD rats treated with imipramine underwent neuropathological evaluation, while rat learning and memory were assessed in the Morris Water Maze. AD patient mortality in the National Alzheimer’s Coordinating Center’s database was analyzed by Kaplan-Meier and Cox proportional hazards models.

**Results:** Imipramine inhibits the apoE-Aβ interaction by binding directly to Aβ monomers. Imipramine reduces Aβ accumulation and neuronal cell loss in AD COs. Imipramine-treated female AD rats had enhanced reference memory and increased hippocampal cellularity and cholinergic neurotransmission. AD patients prescribed imipramine had reduced all-cause mortality over 10 years relative to those prescribed other tricyclic antidepressants (TCA), non-TCA antidepressants, or no antidepressants.

**Conclusions:** The findings support the conclusions that: a) the mechanism by which apoE confers risk for AD is by catalyzing the polymerization of Aβ into neurotoxic species, b) targeting apoE is a promising, underappreciated strategy for the prevention of AD, c) imipramine should be used more frequently to treat depression in AD clinical practice, and d) imipramine should be evaluated in prospective trials, particularly in *APOE4* carriers, as a potential method to improve cognition and reduce mortality in AD.

## BACKGROUND

Inheritance of the ε4 allele of the apolipoprotein E gene (*APOE4*) remains the greatest genetic risk factor for Alzheimer’s disease (AD), and the greatest overall risk factor besides age itself. Despite *APOE4* having an allelic frequency of less than 14% in the general population, approximately two-thirds of individuals diagnosed with AD are *APOE4* carriers. (1, 2) Once thought to be involved in the clearance of amyloid-β (Aβ) peptides from the brain, it is now appreciated that apoE plays numerous important roles in the pathogenesis of AD, including via direct interaction with Aβ, resulting in accelerated oligomer, fibril, and plaque formation.(3-13)

All current FDA-approved disease modifying treatments for AD are monoclonal antibodies that target Aβ and are variably effective at removing amyloid from the brain but also have a high risk for serious adverse events, especially in *APOE4* carriers.(14, 15) Thus, new therapeutic approaches targeting apoE and other disease mechanisms are warranted. Several apoE-targeting therapies have been developed and are currently being evaluated in humans. These include molecules that increase cholesterol transport, enhance Aβ clearance, or genetically manipulate the *APOE* locus to express *APOE2* rather than *APOE4.*(16, 17) *APOE*-targeted therapies have been efficacious in preclinical models, yet whether they will be sufficient to slow or halt a neurodegenerative disease in humans is unclear.

Here we describe an entirely different approach focused on the interaction between apoE and Aβ. We previously developed a high-throughput drug screen to identify small molecule compounds that block the catalytic effect of apoE4 on Aβ polymerization.(18) We screened nearly 600 compounds present in the US NIH Clinical Collection drug library which includes many drugs already approved by the US FDA for various indications. We identified imipramine as the most potent, non-neurotoxic molecule that can cross the blood-brain barrier.(18) Imipramine is a tricyclic antidepressant with norepinephrine and serotonin reuptake inhibitor activity that has seen limited use in the clinic since the development of selective serotonin/norepinephrine reuptake inhibitors (SSRIs/SNRIs).(19, 20) By querying the National Alzheimer’s Coordinating Center (NACC) database, we then revealed that imipramine use to treat depression in AD participants was associated with improved cognitive scores and clinical AD diagnosis.(18) Here, we evaluate the mechanism of action and both preclinical and clinical efficacy of imipramine for the treatment and prevention of AD.

## METHODS

### NMR spectroscopy

NaOH-pretreated Aβ42 (rPeptide) and Aβ40 (Peptide 2.0) were prepared at 20–50 µM, as indicated in the figure legends for each experiment. Imipramine hydrochloride (Sigma-Aldrich) was prepared at 200 µM. We utilized DPBS buffer containing 0.05% NaN_3_ (w/v) and 10% D_2_O (v/v). All NMR measurements were performed at 14.1 T spectrometers (University of Colorado Anschutz and Denver University) equipped with cryoprobes. STD spectroscopy was performed as described previously.(21) Briefly, NMR saturation was applied on-resonance at 0.65 ppm (corresponding to Aβ resonances), and at -40 ppm as a control (off-resonance). A saturation time of two seconds was used and a total of 256 scans per measurement were carried out at 4, 10, and 20 °C. Two-dimensional NOESY measurements were collected with a mixing time of 300 ms. Thirty-two scans, 8192 x136 complex points, and States-TPPI acquisition in the indirect dimension were employed.

^1^H-^15^N HSQC experiments were acquired with a non-uniform sampling (NUS) scheme generated by the NUS@HMS scheme generator employing 1024 complex data points in the direct dimension and 50% sampling of the original 200 complex points in the indirect ^15^N dimension. The spectral widths were 13.7 and 35 ppm for the ^1^H and ^15^N dimensions, respectively, with a relaxation delay of 1.3 s and 32 scans. The data were processed using the iterative soft-thresholding reconstruction method. Data reconstruction of the 2D NUS spectra was performed using the hmsIST software.(22)

All 2D NMR data processing and visualization were conducted using NMRpipe/NMRDraw.(23) Spectra analysis was carried out using the CCPNmr analysis software v2.5.1.(24) One-dimensional STD data was processed in MATLAB.

### hiPSC-derived COs

hiPSC lines HVRDi001-A (57-year-old male) and UCSD241i-APP2-3 (60-year-old female) were received from the Wicell Research Institute and have been described before.(25, 26) hiPSCs were seeded onto Matrigel (Corning) coated polystyrene and maintained in mTeSR Plus Basal Medium (STEMCELL) and then developed into COs using the STEMdiff Cerebral Organoid Kit (STEMCELL) following the manufacturer’s instructions and established protocols(27). Imipramine hydrochloride (Sigma-Aldrich #10899) was dissolved in sterile DPBS and filtered at 0.22 µm, then added at 1 or 10 µM at each media change from 1.5–4 months of age. COs were flash-frozen in liquid N_2_ and stored at -80 °C for immunoblot analyses or immersed in paraformaldehyde (PFA) fixative and stored at room temperature (RT) for IHC analyses.

### TgF344-AD rat imipramine treatment

TgF344-AD rats were received from Taconic Inc. and maintained on a Fischer 344 background, as described previously.(28) Cohorts of six TgF344-AD female rats per group and five TgF344-AD male rats per group were assigned to imipramine and saline groups by forced randomization to balance age and weight, and littermates were split evenly between the groups whenever possible. During the study, one imipramine-treated female rat lost more than 20% of its body weight, a pre-determined humane endpoint, and two male rats, one from the saline group and one from the imipramine group, drowned during the MWM and were removed from the study.

Rats were handled once per day for four days prior to beginning injections to acclimatize the animals to the handler. Imipramine hydrochloride (Sigma-Aldrich #10899) was dissolved at 10 mg/ml in sterile 0.9% saline and filtered at 0.22 µm. Imipramine was prepared freshly each day within one hour of injection. Rats received daily intraperitoneal injections of either 1 µl/g body weight of imipramine (10 mg/kg) or a similar volume (500 µl) of saline. Left and right sides were alternated each day for injections to minimize pain. Rats were weighed at least every four days, and the injection volume of imipramine was updated accordingly. Every injection was performed by the same animal handler within the same two-hour span each day. Eighteen days after initiating injections, rats were tested for hippocampal-based learning and memory in the MWM. Injections were performed on each day of the MWM and the final injection was performed approximately 12 h prior to sacrifice. The brains were collected, and the right hemispheres were immersed in PFA fixative for IHC analyses while the left hemispheres were divided into cortex, hippocampus, and hindbrain pieces and stored at -80 °C. Blood was collected in heparinized tubes by intracardiac puncture, and plasma was isolated by centrifugation and stored at -80 °C.

### Morris water maze (MWM)

A 170 cm diameter round metal tank was filled with room-temperature water, and black paint was added to provide contrast to the rats’ white coats for video recording and analysis using Ethovision XT. A hidden platform was submerged approximately 1 cm below the surface of the water in the middle of the southeast quadrant of the tank. The tank was surrounded by black and white curtains containing designs that provided visual cues. Rats were habituated to the behavioral testing room for 30 min prior to each testing session. On the first day, rats were habituated to the tank environment by placing them on the platform for 30 s prior to the first training trial. Rats were tested in four training trials per day by placing them in either the west, southwest, north, or northeast area of the tank facing the wall and allowing them to swim until they found the platform, or for 120 s after which they were gently guided to the platform. The same investigator who performed injections also handled the animals during behavioral testing. A second investigator blinded to the treatment group timed the latency to find the platform. Rats were dried and rested for five minutes in a heated recovery cage before the next trial. The release quadrant was alternated for each trial and a different order of release quadrants was used each day. On day five, the platform was removed, and rats were tested in a single 120 s probe trial after release in the northwest quadrant. Rats were humanely euthanized immediately following the probe trial.

### Immunoblotting

COs were thawed and immersed in RIPA buffer (Fisher Scientific) containing 1X protease and phosphatase inhibitor cocktail (Thermo Scientific). COs were lysed by probe sonicator for 10 s and then incubated on ice for 30 min. Rat cortical tissue was thawed on ice, and DPBS containing 1X protease and phosphatase inhibitor cocktail was added to prepare a 25% (w/v) homogenate. Ceramic beads were added, and tissue was homogenized using a bead beater for 30 s. CO and rat cortical protein lysates were centrifuged at 20,000 x g for 20 min at 4 °C to remove tissue debris, and total protein was assessed by BCA assay (Thermo Scientific). Total protein concentrations were normalized, and the samples were then combined with NuPAGE LDS sample buffer and 1X reducing agent (Invitrogen) and heated at 95 °C for 10 min. Two micrograms of CO protein or 20 µg of rat cortical protein was loaded per well in a 4–12% polyacrylamide Bis-Tris gel (Invitrogen), and electrophoresis was performed at 200 V for 45 min in MES buffer. Proteins were transferred to a PVDF membrane using an iBlot™ 2 Gel Transfer Device (Thermo Scientific) at 20 V for 7 min. PVDF membranes were incubated in blocking buffer containing 3% BSA in TBS buffer containing 0.01% Tween-20 (TBST) for 2 h at RT. Membranes were then incubated with primary antibody labeling both APP and Aβ (6E10, Biolegend, 1:1,000) in blocking buffer overnight at 4 °C. Membranes were washed three times in TBST and then incubated with HRP-conjugated goat anti-mouse IgG secondary antibody (Jackson Immunolabs, 1:5,000) in blocking buffer for 1 h at RT. Membranes were washed in TBST, incubated with SuperSignal West Pico Plus ECL substrate (Thermo Fisher) for 90 s, and developed on an iBright FL 1500 Imaging System (Thermo Fisher). Membranes were stripped for 10 min in 1X ReBlot Plus Mild Stripping Solution (Millipore) and re-probed with a primary antibody against β-actin (Santa Cruz Biotech, 1:2,000) and other steps were identical. Band intensities of APP (∼100 kDa), Aβ (∼4 kDa), and β-actin (∼42 kDa) were quantified using ImageJ 1.54.

### Immunohistochemical (IHC) analysis

COs were fixed in 4% PFA in DPBS overnight at 4 °C and rat hemi-brains were fixed for 72 h at RT. Tissues were then taken through a stepwise concentration gradient of sucrose in DPBS: 6.25% overnight, 12.5% overnight, and then 25% overnight at RT. Fixed COs and rat hemi-brains were embedded in Tissue-Tek OCT Compound (Sakura), frozen by immersing the molds in liquid N_2_-cooled isopentane, cryosectioned at 20 µm thickness, and stored at -80 °C.

For immunostaining, slides were washed three times in DPBS to dissolve OCT and then incubated for 1 h in permeabilization/blocking buffer [0.25% Triton X-100 and 3% bovine serum albumin (BSA) in DPBS]. Human CO sections were incubated with primary antibodies against microtubule-associated protein 2 (MAP2; PhosphoSolutions, 1:1,000) and glial fibrillary acidic protein (GFAP; Millipore, 1:500), and rat brain sections were incubated with primary antibodies against tau paired helical filaments (PHF1; Dr. Peter Davies, 1:50), or neuronal nuclei (NeuN; Abcam, 1:500), all diluted in staining buffer (0.1% Triton X-100 and 3% BSA in DPBS) and incubated in the dark overnight at 4 °C. Slides were washed in DPBS and sections were covered with AlexaFluor Plus-conjugated goat anti-mouse IgG or goat anti-rabbit IgG secondary antibodies (Invitrogen, 1:500) in staining buffer and incubated in the dark for 2 h at RT. Sections were counterstained with AmyloGlo (Biosensis, 1:200) and Hoechst 33342 or DRAQ5 (EBioscience, 1:2,000) in DPBS for 10 min at RT, washed in DPBS, and coverslips were added with Prolong Gold antifade mountant (Invitrogen). Images were acquired on an Olympus IX83 inverted fluorescence microscope and analyzed using Cell Sens v1.12 software (Olympus).

### Liquid chromatography-mass spectrometry (LC-MS)

The concentrations of imipramine in plasmas and cortical brain lysates of TgF344-AD rats at sacrifice were determined by liquid chromatography-mass spectrometry (LC-MS)-based metabolomics. Imipramine was extracted using a modified protein crash protocol described previously.(29-31) Briefly, 20 µL of plasma or brain lysate was combined with 180 µL of cold 5/3/2 MeOH/ACN/H_2_O (v/v/v) and vortexed at 4 °C for 30 min. Samples were centrifuged at 18,213 x g for 10 min at 4 °C and the supernatant was transferred to HPLC autosampler vials. Portions of the extracts from each sample were also combined and injected throughout the run for quality control. Two calibration curves were prepared by spiking imipramine into plasma or brain lysate from untreated TgF344-AD rats, with calibrants including 1, 5, 50 and 100 ng/mL for plasma, and 1, 5, 10, 50, 75 and 100 ng/mL for brain lysate. The final concentrations of imipramine were determined by weighting the curves linearly 1/x^2^.

Analyses were performed using a modified gradient optimized for high-throughput quantitative metabolomics, as described previously.(29) Briefly, a Vanquish UHPLC system (Thermo Fisher) coupled to a QExactive mass spectrometer (Thermo Fisher) was used. Imipramine extracts were resolved with an ACQUITY UPLC BEH C18 column (Waters, 2.1 x 100 mm, 1.7 µm particle size) held at 45 °C. For positive ion mode analysis, mobile phases (A) 0.1% FA in H_2_O, and (B) 0.1% FA in ACN were used in a chromatographic gradient: 0% B at 0 min, 0% B at 0.5 min, 100% B at 1.1 min, 100% B at 2.75min, 0% B at 3min, and 0% B at 5 min, with a constant flow rate of 0.45 mL/min. The mass spectrometer scanned in Full MS mode from 65 to 975 m/z at 120,000 resolution, with 4 kV spray voltage, 45 sheath gas, 25 auxiliary gas. Calibrations were performed prior to analysis using the PierceTM Positive and Negative Ion Calibration Solutions (Thermo Fisher). Acquired data was converted from raw to mzXML file format using RawConverter. Analysis was done using El MAVEN, an open-source software program for chromatographic identification. Samples were analyzed in randomized order with a technical mixture injected interspersed throughout the run to qualify instrument performance.

### Enzyme-linked immunosorbent assay (ELISA)

Rat cortical tissue homogenates (25% w/v) were thawed on ice, and formic acid (FA) extraction was performed as previously described.(32) Briefly, 25 µL homogenate was combined with 50 µl formic acid (FA) in a microcentrifuge tube. The samples were vortexed, sonicated for 20 min at 37 °C in a water-bath sonicator, and centrifuged at 26,000 × g for 30 min. Fifty µl of supernatant were moved to a low-binding tube and neutralized with 950 µL of neutralization buffer (1 M Tris base and 500 mM dibasic sodium phosphate). FA-extracted samples were diluted 32-fold in assay buffer and analyzed using the Human Aβ42 ELISA kit (R&D Systems) according to the manufacturer’s instructions. Samples were analyzed on a Synergy H4 Hybrid plate reader (Biotek).

### AChE and BChE activity assays

Rat plasma samples were thawed on ice and cholinesterase activities were measured using AChE and BChE assays (Thermo Fisher) according to the manufacturer’s instructions. Briefly, for the AChE assay both male and female samples were diluted 500-fold in the supplied assay buffer. For the BChE assay, male samples were diluted 30-fold and female samples were diluted 60-fold. Samples were analyzed on a Synergy H4 Hybrid plate reader (Biotek).

### NACC data analysis

The NACC uniform dataset v3 was received on October 27, 2025, and contained standardized longitudinal clinical data on subjects seen at Alzheimer’s Disease Research Centers (ADRCs) beginning in September 2005 through the September 2025 data freeze.(33) Subjects who reported taking the medications studied here were identified by searching the ‘DRUGS’ column, as described previously.(18) For Kaplan-Meier analyses, medications were arranged in a hierarchy (top to bottom): imipramine, TCA medications other than imipramine, non-TCA antidepressants, and no anti-depressants. Each patient was assigned to a medication group depending on the highest tier medication, and their exposure time was started seven months after their first observation to ensure that the medications had time to take effect. Thus, only subjects who had a record beyond seven months of exposure time were considered. The outcome was time to mortality within ten years after start, and time for patients without a recorded death were censored at the last time of observation or at ten years. The numbers of participants at risk and censored are provided in **Data S1**. Kaplan-Meier plots were created for the medications, and the medication groups were compared using the Peto-Peto test. Survival probabilities were estimated for times of 2.5, 5.0, 7.5, and 10.0 years, along with log-log pointwise 95% confidence intervals. The differences between medication groups were summarized by fitting a Cox proportional hazards model and reporting and testing the hazard ratios among groups. All tests were two-sided univariate α = 0.05 unless otherwise stated.

Protective omnibus tests were performed. Statistical computations were performed in SAS 9.4, with GraphPad PRISM 11 and STATA 17 used for graphical representations.

### Statistical analysis

Analysis of variance (ANOVA) and t-tests were performed using GraphPad Prism 11. Following ANOVA, comparisons between multiple groups were done by post-hoc testing using the Holm-Šidák method and a *P* < 0.05 was considered statistically significant. All statistical tests were two-sided. All measurements were taken from distinct samples.

## RESULTS

### Imipramine binds to Aβ42 monomers

The interactions between imipramine and Aβ were probed using several nuclear magnetic resonance (NMR) spectroscopy methods. First, 2D ^1^H-^1^H Nuclear Overhauser Effect Spectroscopy (NOESY) was utilized. In a classic 2D NOESY spectrum, binding between the ligand and protein is detected as a change of sign in the cross-peaks of the ligand. A free ligand behaves as a small molecule in the fast-tumbling regime, which results in a difference in the phase between diagonal and cross peaks (i.e., negative cross-peaks, which are of opposite sign to the diagonal). On the other hand, a ligand bound to a macromolecule such as Aβ behaves as a large molecule due to the slower tumbling and hence cross and diagonal peaks for this ligand will have the same phase/sign.(34, 35) This was indeed the case for free imipramine (**Fig. 1a**), and for imipramine bound to Aβ42 monomer (**Fig. 1b**). Additionally, there was a clear broadening of the cross-peaks for imipramine in the presence of Aβ42 (**Fig. S1**), serving as additional confirmation of binding and an increase in the tumbling time. Interestingly, no characteristic features of binding (change in sign or broadening of cross-peaks) were observed in the NOESY spectrum of impramine with Aβ40 (**Fig. S2**), thus its interaction with Aβ42 appears to be specific.

**Figure 1.**
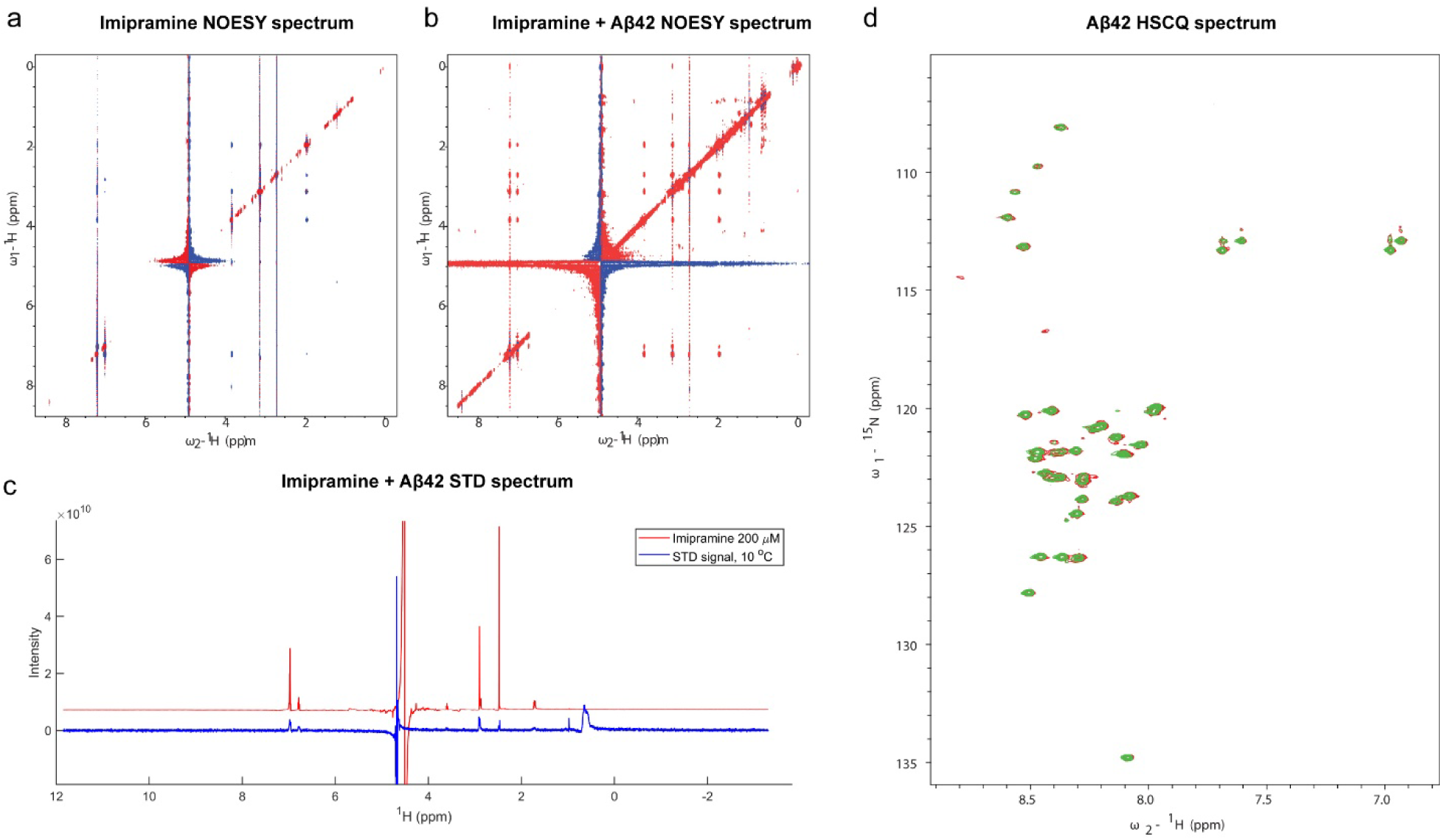
Imipramine binding to Aβ42 revealed by NMR. a,b) 1H-1H NOESY spectra of (a) 200 µM imipramine alone, and (b) in the presence of 25 µM Aβ42 monomers at 10 °C. The mixing times were 300 ms. Negative cross-peaks are shown in red and positive cross-peaks are shown in blue. Binding is observed in (b) by the change of sign of the cross-peak of imipramine upon binding, as well as the change in widths of the peaks. c) 1H NMR STD spectra for 20 µM Aβ42 and 200 µM imipramine at 10 °C. Intensity in arbitrary units versus chemical shift. Red line: imipramine 1H NMR spectrum in isolation, blue line: STD signal difference of the mixture, scaled by a factor of 23 relative to the red trace. d) 1H-15N HSQC spectrum of 50 µM Aβ42 at 10 °C. No chemical shift perturbation or changes in intensities were observed in the presence of 200 µM imipramine.

Next, Saturation Transfer Difference (STD) spectroscopy was utilized which is based on the transfer of saturation of resonances from the macromolecule to the ligand. During STD spectroscopy, low-power irradiation is applied to the region of the spectrum containing some isolated protein signal, which spreads throughout the entire macromolecule via spin diffusion and results in magnetization transfer to the ligand if it is transiently bound to the macromolecule(21). The experiment was performed twice: once with the Gaussian saturation pulse applied on resonance to target the Aβ42 monomer protein methyl resonances, and again off-resonance as a control. The two spectra were subtracted, revealing the STD effect. In the spectrum of free imipramine, no STD signal difference was observed at concentrations below 400 µM, as expected. In the presence of Aβ42 monomer, however, non-zero STD signal differences were observed for imipramine, which indicates binding and distance proximity (**Fig. 1c**), and this was confirmed at multiple temperatures (**Fig. S2**).

Finally, heteronuclear single-quantum correlation (HSQC) NMR spectra were collected for ^15^N-labeled Aβ42 monomers alone and in the presence of imipramine. For a structured protein, changes in the backbone resonance positions or intensities are expected upon ligand binding for the sites affected by the ligand. However, no chemical shift changes were observed (**Fig. 1d**). Because Aβ42 monomer is disordered, the subtle changes in the conformational ensemble are not visible with this technique, suggesting that binding with imipramine may be more pronounced at the side-chain sites.

### Imipramine treatment reduces Aβ levels in AD cerebral organoids

The effects of imipramine on AD neuropathologies were next evaluated in hiPSC-derived cerebral organoids (COs). hiPSC lines derived from two participants with autosomal dominant AD: one female with a duplication of the amyloid precursor protein (*APP*) gene (APPdup), and one male with the APP London mutation (V717I), were developed into COs. Both mutations result in increased production and accumulation of amyloid-β (Aβ) peptides, leading to robust brain amyloid and early-onset AD dementia.(25, 26, 36-38) AD COs were treated with 1 or 10 µM imipramine at 1.5–4 months of age, beginning prior to the development of Aβ neuropathology in AD COs.(39-41) At collection, there was no difference in the gross size of the AD COs treated with imipramine compared to control AD COs (**Fig. 2a,b**). Although their APP levels were similar (**Fig. 2c**), AD COs treated with 1 µM or 10 µM imipramine had significantly lower levels of monomeric and oligomeric Aβ relative to control AD COs (**Fig. 2d**), as assessed by immunoblotting. AD COs treated with 1 or 10 µM imipramine also had greater GFAP^+^ astrocytic and MAP2^+^ neuronal compositions relative to control AD COs (**Fig. 2e–g**), likely indicating greater astrocytic reactivity to Aβ and reduced neurodegeneration with imipramine treatment.

**Figure 2.**
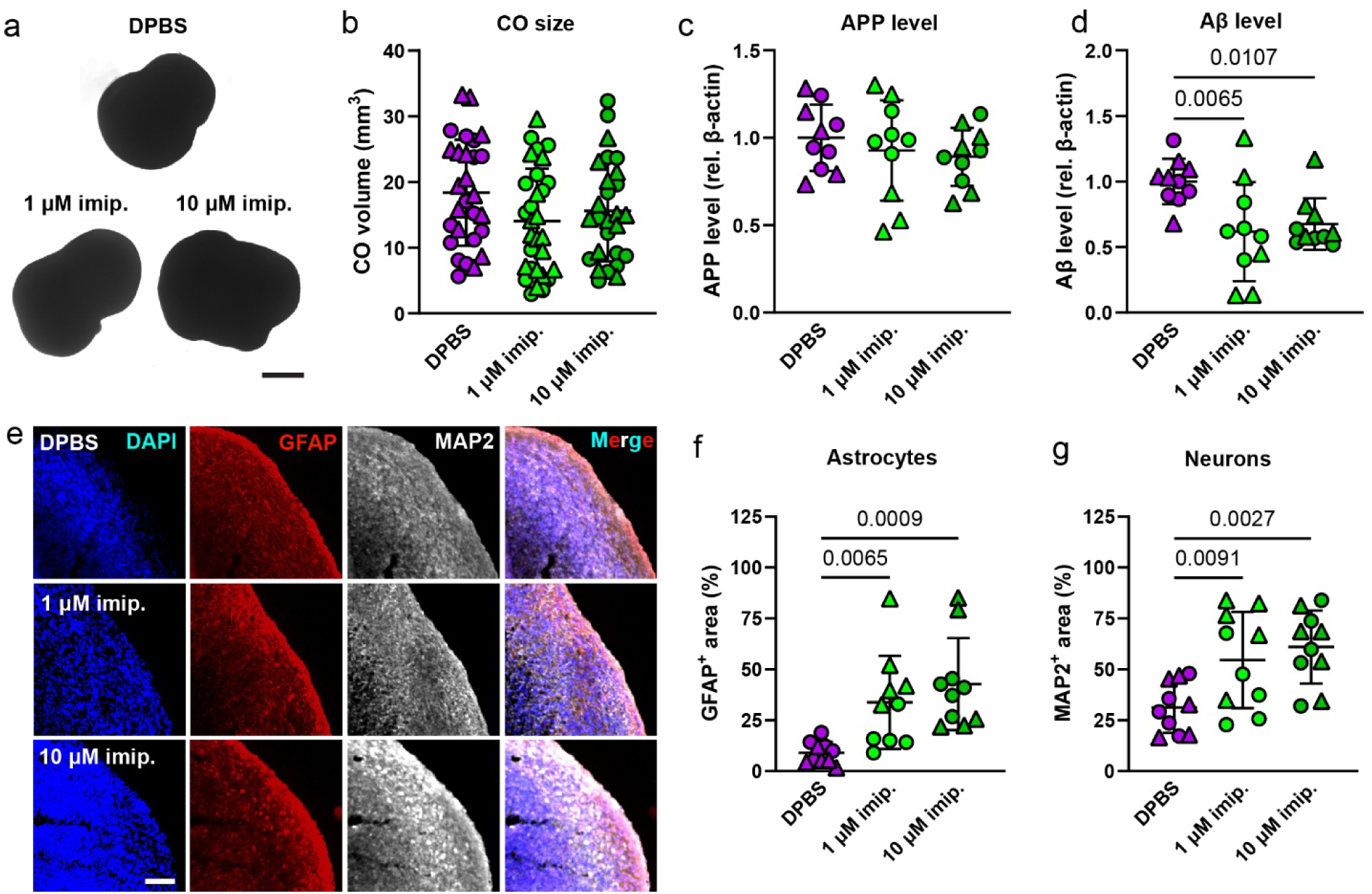
Imipramine reduces Aβ levels in hiPSC-derived AD COs. a) Representative light microscopy images of hiPSC-derived autosomal dominant AD COs treated with 1 or 10 µM imipramine or Dulbecco’s Phosphate Buffered Saline (DPBS) as a negative control. Scale bar = 1 mm. b) Volumes of AD COs, estimated from measured perimeters and assuming a spherical shape. hiPSCs were derived from one female with APPdup (circles) and one male with APPV717I (triangles). Each symbol represents an individual CO. c) APP levels in AD CO tissue lysates measured by immunoblotting, relative to β-actin levels. d) Aβ levels in AD CO tissue lysates measured by immunoblotting, relative to β-actin levels. One-way ANOVA with Holm-Sidak post-hoc testing. e) Representative immunohistochemistry (IHC) images of AD COs derived from a male with APPV717I, immunolabeled for glial fibrillary acidic protein (GFAP) and microtubule associated protein 2 (MAP2) and counterstained with DAPI. Scale bar = 100 µm. f) Quantification of GFAP^+^ astrocytic area fraction in AD COs. One-way ANOVA with Holm-Sidak post-hoc testing. g) Quantification of MAP2^+^ neuronal area fraction in AD COs. One-way ANOVA with Holm-Sidak post-hoc testing.

### Imipramine treatment enhances reference memory in female AD rats

TgF344-AD rats express the human APP gene harboring the Swedish (K670N/M671L) mutation and the human PSEN1 gene with the Δ exon 9 mutation and develop the full range of brain neuropathology and cognitive changes associated with AD in humans.(28, 42, 43) Both male and female rats, approximately 16–20 months of age, were used in the study. Although the male rats were generally older (t = 6.434, *P* < 0.0001) and heavier (t = 5.181, *P* < 0.0001) than the female rats, there was no significant difference in age (F = 0.1831, *P* = 0.6748) or body weight (F = 0.1238, P = 0.7298) between the saline and impramine groups within each sex at treatment initiation (**Table S1**).

After treatment initiation, there was a significant treatment × time interaction (**Fig. 3a,b**; F = 49.68, *P* < 0.0001) and no significant treatment × sex interaction (**Fig. 3a,b**; F = 0.07368, *P* = 0.7898), indicating that both male and female imipramine-treated rats had greater weight loss than saline-treated rats, as has been reported before in wild-type rats.(44) Although the imipramine concentrations at sacrifice were not significantly different between female and male rats in either their brains (**Fig. 3c**) or their plasmas (**Fig. 3d**), there was a significantly higher ratio of imipramine in the brain relative to the plasma in female rats compared to male rats (**Fig. 3e**), possibly due to reduced brain penetrance of imipramine in males or reduced brain clearance of imipramine in females.

**Figure 3.**
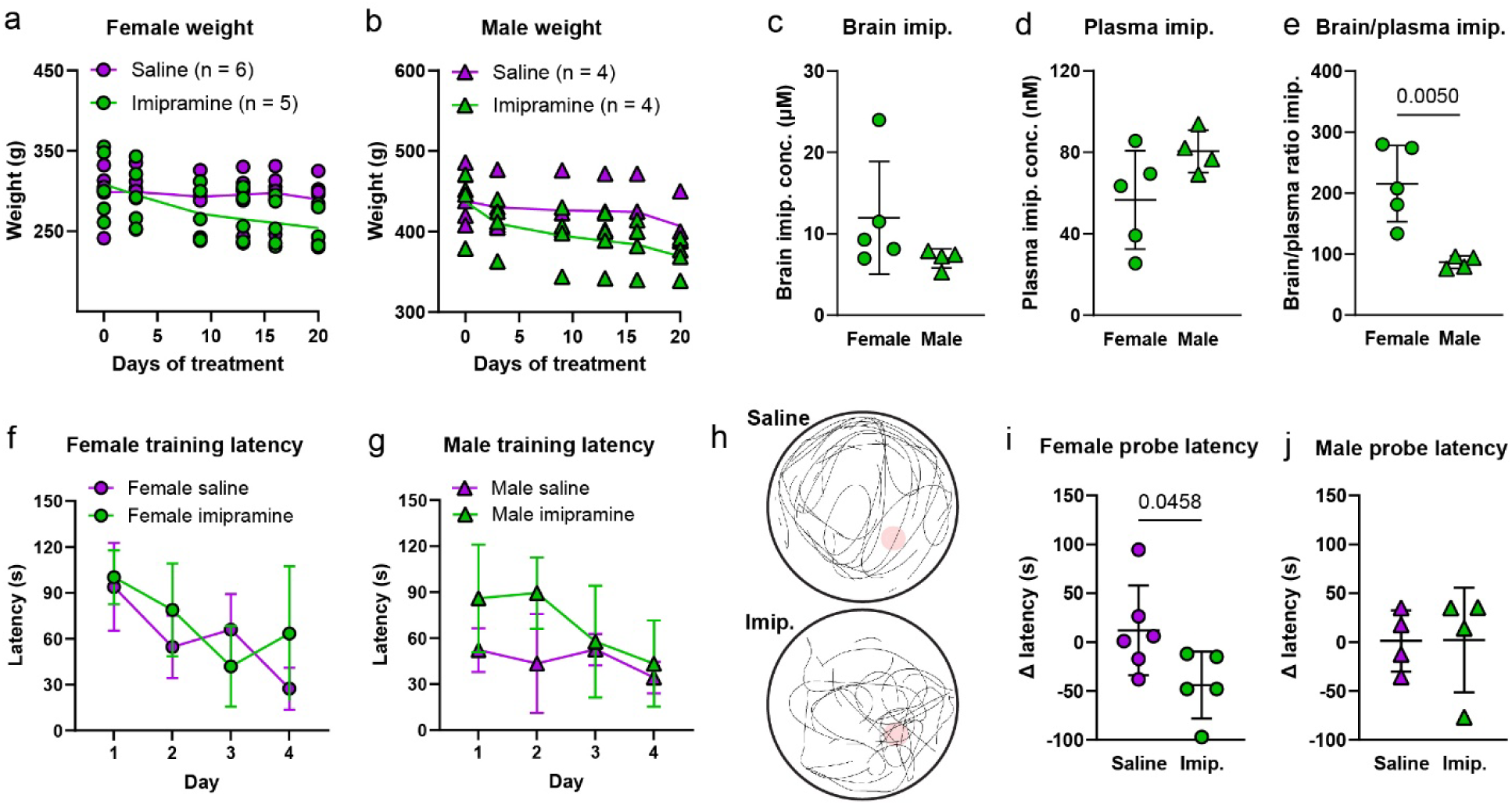
TgF344-AD rat imipramine pharmacokinetics and Morris water maze (MWM) performance. a,b) Weights of (a) female and (b) male TgF344-AD rats over 20 days of daily treatment with imipramine (10 mg/kg i.p.) or a similar volume of saline. Each symbol represents an individual rat, and lines indicate group means. c,d) Concentrations of imipramine in (c) cortical brain lysates and (d) plasmas of TgF344-AD rats at sacrifice, approximately 12 h following the final injection, determined by liquid chromatography-mass spectrometry (LC-MS)-based metabolomics. Welch’s unpaired t-test. Lines indicate group means ± SD. e) The brain-to-plasma ratios of imipramine concentrations were calculated for each rat. Welch’s unpaired t-test. f,g) Average latency to the platform of (f) female and (g) male TgF344-AD rats during the four training days of the MWM. h) Representative traces of the paths (thin black lines) of saline- and imipramine-treated TgF344-AD female rats during the MWM probe trial. The outer circle (thick black line) indicates the boundary of the tank with the platform zone location (red circle) shown in the southeast quadrant. i,j) Change in latency to the platform zone between the final training and probe trials of (i) female and (j) male TgF344-AD rats. Welch’s unpaired t-test.

Aged TgF344-AD rats (15- to 16-month-old) have been previously reported to exhibit impaired hippocampal-based spatial learning and memory in the Barnes maze and in the Morris water maze (MWM).(28, 42) Here we performed the MWM, and on the first day of training there was no significant effect of imipramine treatment on the distance swum, average swimming velocity, or latency to reach the platform relative to saline treatment (**Fig. S4a**). However, there was a statistically significant main effect of sex on the distance swum (**Fig. S4b**; F = 5.409, *P* = 0.0345) and the latency to reach the platform (**Fig. S4c**; F = 5.602, *P* = 0.0318) with male rats swimming shorter distances and reaching the platform faster than female rats. These data reveal a sex-specific difference in TgF344-AD rat hippocampal-based learning not identified in prior studies, and thus the effects of imipramine treatment on male and female rats were analyzed separately.

Imipramine treatment did not significantly affect learning or working memory of either female (**Fig. 3f**; F = 0.8014, *P* = 0.3940) or male rats (**Fig. 3g**; F = 3.241, *P* = 0.1219) relative to saline treatment during the four-day MWM training phase as measured by their latency to reach the platform. In the probe trial on day 5, reference memory of each rat was assessed by their change in latency to reach the platform relative to day 4 of their training. Female imipramine-treated rats reached the platform zone quicker and spent more time in the target quadrant (**Fig. 3h**), thus exhibiting better reference memory relative to female saline-treated rats (**Fig. 3i**), while no effect of imipramine treatment on reference memory was observed in male rats (**Fig. 3j**). Consistent with the hypothesis that imipramine had a beneficial effect on reference memory in female rats but not in male rats, imipramine concentration in the brain at sacrifice was correlated with better performance in females (**Fig. S5a**, R^2^ = 0.8034; *P* = 0.0395) and worse performance in males (**Fig. S5b**, R^2^ = 0.9229; *P* = 0.0393).

### Imipramine treatment reduces neurodegeneration in the brains of AD rats

TgF344-AD rats have been reported to accumulate both soluble and detergent-insoluble Aβ42 in the hippocampus and cerebral cortex in an age-dependent manner.(28) In the present study, despite similar levels of APP (**Fig. 4a, F** = 1.610; *P* = 0.2238), female TgF344-AD rats had a notable trend towards higher baseline levels of formic acid (FA)-extracted Aβ in cortical lysates compared to male rats (**Fig. 4b, F** = 4.264; *P* = 0.0567). Likely due to their higher baseline levels, the reduction in cortical Aβ levels with imipramine treatment relative to saline treatment was significant in female rats (**Fig. 4b**), but not in male TgF344-AD rats.

**Figure 4.**
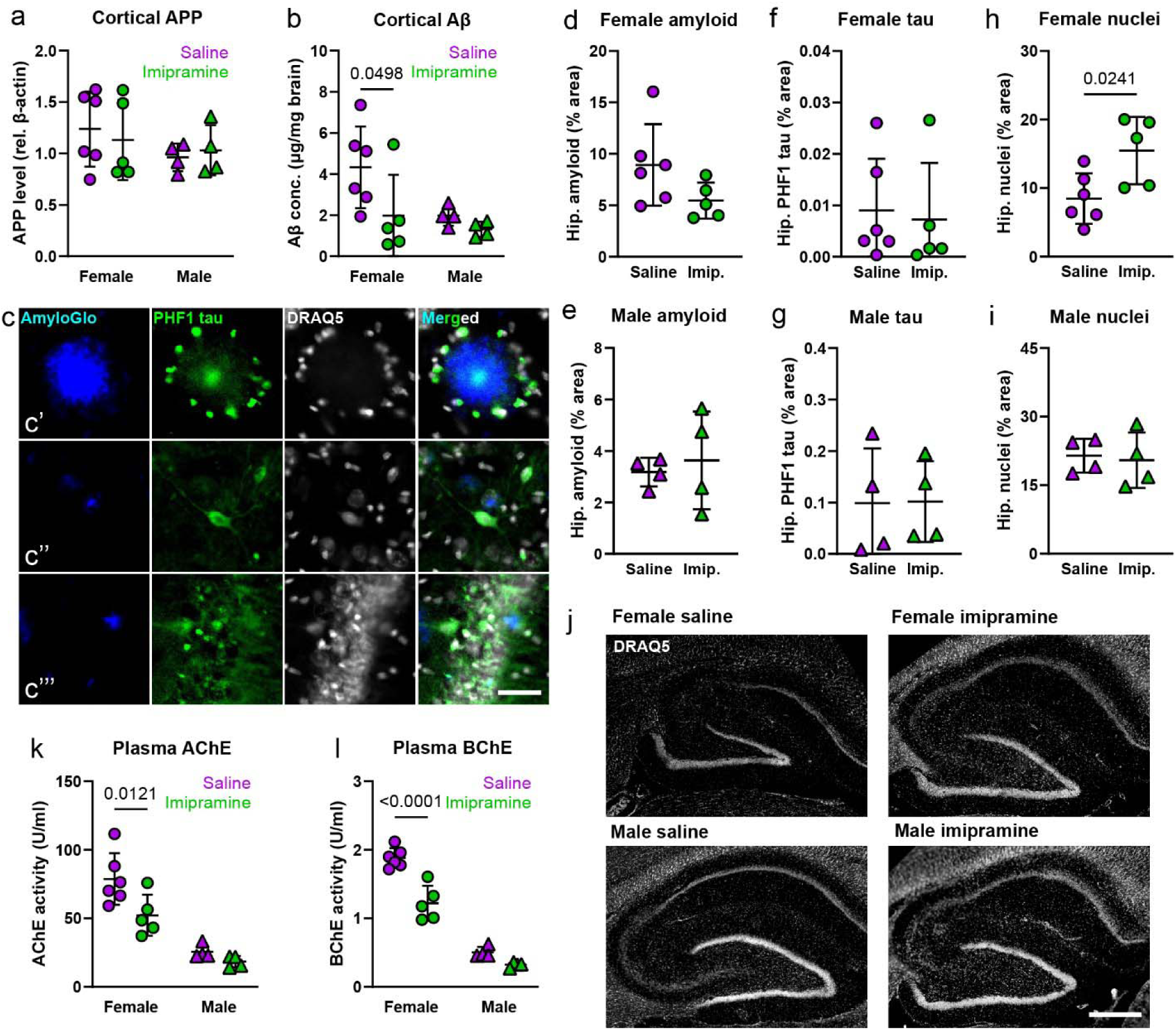
Effects of imipramine on Aβ, neurodegeneration, and cholinesterase levels in TgF344-AD rats. a) APP levels in TgF344-AD rat cortical tissue lysates measured by immunoblotting, relative to β-actin levels. Each symbol represents an individual rat. Lines indicate group means ± SD. b) Aβ concentration in formic acid-extracted TgF344-AD cortical tissue lysates measured by enzyme-linked immunosorbent assay (ELISA) and normalized to total protein level. Two-way ANOVA with Holm-Sidak post-hoc testing. c) Representative IHC images of a saline-treated female TgF344-AD rat brain showing a cortical amyloid plaque (c’), and intraneuronal Aβ aggregation in the cortex (c’’) and hippocampus (c’’’), labeled for amyloid (AmyloGlo), tau paired helical filaments (PHF1), and cell nuclei (DRAQ5). Scale bar = 20 µm. d,e) Quantification of AmyloGlo^+^ amyloid area fraction in the hippocampi of saline- and imipramine-treated (d) female and (e) male TgF344-AD rats. Welch’s unpaired t-test. f,g) Quantification of PHF1^+^ tau area fraction in the hippocampi of saline- and imipramine-treated (f) female and (g) male TgF344-AD rats. Welch’s unpaired t-test. h,i) Quantification of DRAQ5^+^ nuclei area fraction in the hippocampi of saline- and imipramine-treated (h) female and (i) male TgF344-AD rats. Welch’s unpaired t-test. j) Representative IHC images of the hippocampi of saline- and imipramine-treated female and male TgF344-AD rats labeled for cell nuclei (DRAQ5). Scale bar = 500 µm. k) Acetycholinesterase (AChE) activity levels in the blood plasmas of saline- and imipramine-treated TgF344-AD rats. Two-way ANOVA with Holm-Sidak post-hoc testing. l) Butyrylcholinesterase (BChE) activity levels in the blood plasmas of saline- and imipramine-treated TgF344-AD rats. Two-way ANOVA with Holm-Sidak post-hoc testing.

TgF344-AD rats have also been found to develop intraneuronal Aβ in the cerebral cortex and hippocampus identified by electron microscopy and co-immunolabeling.(28) We reported previously that imipramine dose-dependently reduces intraneuronal Aβ accumulation *in vitro* in primary neurons from TgF344-AD rats.(18) In the present study, in addition to extracellular dense-core amyloid plaques, smaller Aβ aggregates closely associated with cellular nuclei were also observed in the saline-treated TgF344-AD rat cortex and hippocampus (**Fig. 4c**), potentially revealing intraneuronal accumulation of Aβ.

Consistent with the sex-dependent differences in Aβ levels in the brains of TgF344-AD rats, sex-dependent differences in the amyloid and tau neuropathological loads were also observed. Hippocampal amyloid neuropathology was greater in female rats relative to male rats (F = 9.699; *P* = 0.0071) while, conversely, tau neuropathology was greater in male rats relative to female rats (F = 11.06; *P* = 0.0046). Aged TgF344-AD rats have been reported to lose approximately 40% of hippocampal neurons relative to wild-type rats.(28, 45) A sex-dependent difference in hippocampal cellularity was observed with male rats having a greater number of cell nuclei relative to female rats (F = 17.78; *P* = 0.0007), as well as a greater number of NeuN^+^ neuronal nuclei (F = 20.03; *P* = 0.0004). A similar sexual dimorphism has been reported previously in the brains of aging rhesus macaques.(46) Due to these sex-dependent differences, the effects of imipramine on amyloid, tau, and neurodegeneration were assessed separately in male and female rats.

No significant difference in amyloid (**Fig. 4d,e**) or tau neuropathology (**Fig. 4f,g**) was observed in the hippocampi of either female or male TgF344-AD rats treated with imipramine relative to saline-treated rats. However, a trend towards reduced hippocampal amyloid in impramine-treated female rats was observed (**Fig. 4d**, t = 1.809, *P* = 0.1040), likely due to reduced Aβ concentrations in their brains relative to saline-treated rats. Imipramine-treated female rats also had increased cellular nuclei in their hippocampi relative to saline-treated rats, while imipramine-treated male rats did not (**Fig. 4h–j**). Finally, there was a trend towards increased NeuN^+^ hippocampal nuclei in imipramine-treated female rats relative to saline-treated female rats (**Fig. S6a**, t = 1.992, *P* = 0.0775), but not in imipramine-treated male rats (**Fig. S6b**, t = 0.6114, *P* = 0.5634). These data indicate that imipramine-treated female TgF344-AD rats benefitted from reduced neurodegeneration, likely due to reduced brain Aβ levels, which in part explains their enhanced MWM performance.

### Imipramine treatment reduces plasma cholinesterase activity in AD rats

Aged TgF344-AD rats display signs of cholinergic dysfunction, evidenced in part by reduced hippocampal acetylcholine levels relative to wild type rats.(28, 42, 47) TgF344-AD female rats were found to have significantly higher activity levels of both acetylcholinesterase (AChE) and butyrylcholinesterase (BChE) in their blood plasma compared to male rats (AChE: F = 46.50; *P* < 0.0001, BChE: F = 227.9; *P* < 0.0001), as has been reported before in wild type rats.(48) There was a significant reduction in the activity levels of both AChE and BChE in the plasma of imipramine-treated female rats relative to saline-treated rats (**Fig. 4k,l**). Reduced cholinesterase activity results in enhanced choline-dependent neurotransmission, suggesting that imipramine may protect cholinergic neurons or boost cholinergic function in the brains of TgF344-AD rats.

### Imipramine treatment extends the survival of AD patients

In retrospective analyses of the National Alzheimer’s Coordinating Center (NACC) clinical dataset, we previously reported that participants who were prescribed imipramine had significantly better cognition as measured by the MMSE and a subsequent reduction in diagnosis of AD relative to participants who were prescribed any other antidepressant medication(18). Here, we assessed whether the same participants also survived longer, potentially due to a reduced risk for AD-related mortality. Participants were identified who had been prescribed imipramine, any other tricyclic antidepressant (TCA), any non-tricyclic antidepressant medication (non-TCA), primarily SSRI/SNRIs, or those who had no recorded prescription of an antidepressant medication (**Table S2**).

Kaplan-Meier (KM) survival analysis revealed that participants prescribed imipramine had significantly higher rates of survival over 10 years relative to participants prescribed any other TCA (*P* = 0.0450), any non-TCA (*P* = 0.0012), or no antidepressants (**Fig. 5a**; *P* = 0.0263). The Peto-Peto test was used because it weights by an estimated pooled survival function, thus weighting earlier times more than later times, thereby increasing the power for detecting differences from the imipramine group, which only had 16 deaths. Participants prescribed imipramine had an estimated 81% probability of survival at 5 years and 66% probability of survival at 7.5 years, compared to 50% probability of survival at 5 years and 33% probability of survival at 7.5 years for the participants prescribed a non-TCA antidepressant (**Table 1**). The log rank test, which weights all times equally, also had an omnibus test *P* value that was statistically significant (**Table S3**; *P* < 0.0001). However, the Cox supremum test, as well as the KM survival plot and the accompanying Epanechnikov kernel-smoothed hazards plot (**Fig. 5b**), indicated some deviation from the proportional hazards assumption, particularly for the Other TCA group. As a result, the imipramine group was only statistically significantly different from the non-TCA group (**Table S3**; P = 0.0094) by the log rank test.

**Figure 5.**
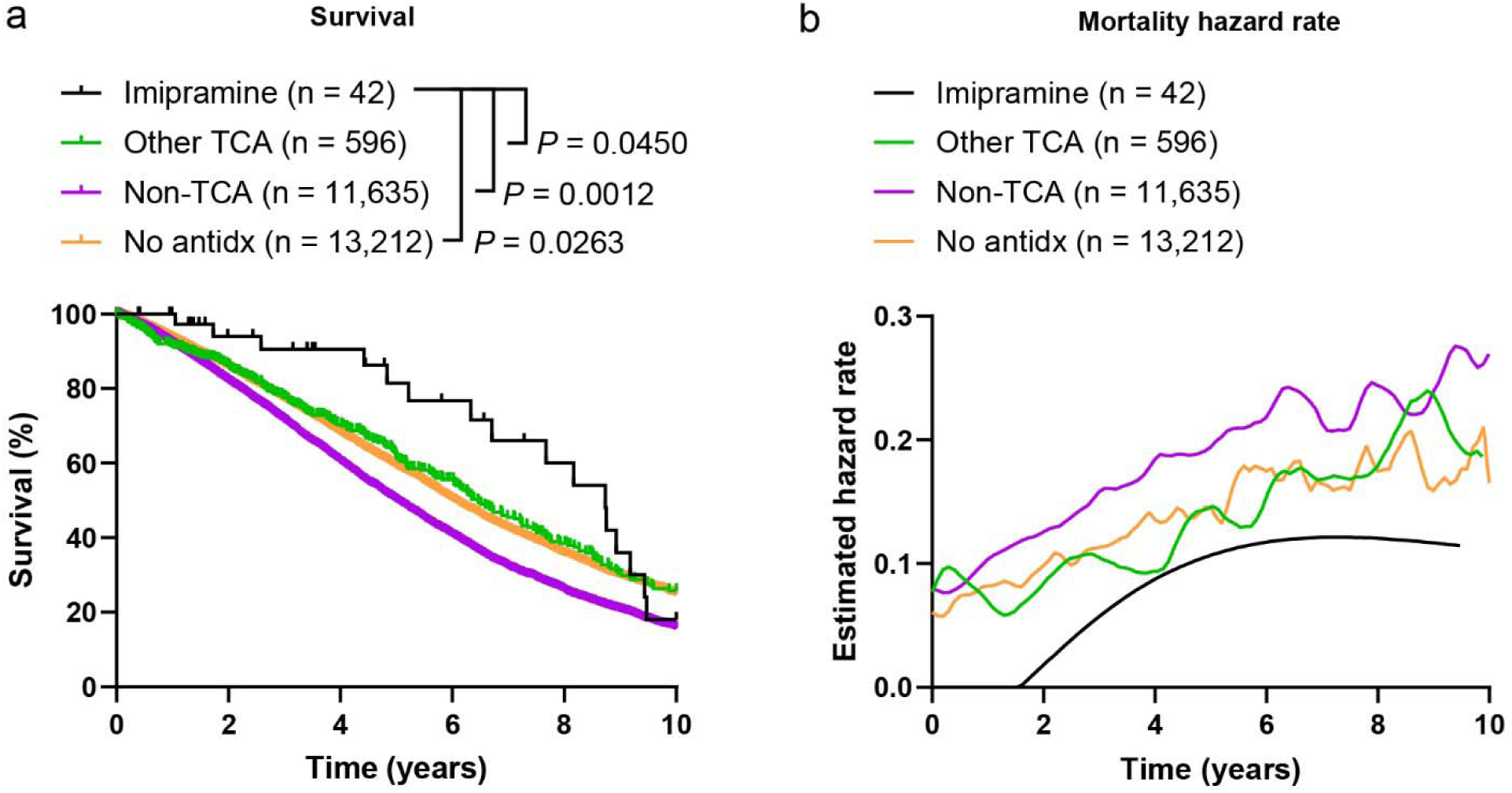
NACC survival and mortality hazard analyses. NACC participants seen at Alzheimer’s Disease Research Centers who reported being prescribed imipramine, any other tricyclic antidepressant (TCA), any non-TCA antidepressant, or no antidepressants (antidx) were identified. a) Kaplan-Meier (KM) survival analysis was performed. Exposure time started 7 months after first observation to allow the medications time to take effect and survival over 10 years was compared by Peto-Peto test. Numbers of participants at risk and censored are provided in **Data S1**. b) The Epanechnikov kernel-smoothed hazard functions for each group, derived from the KM analysis, were plotted.

**Table 1.** Temporal Kaplan-Meier survival probability estimates. Confidence intervals are point-wise and were calculated using the log-log transform.

|  | Kaplan-Meier survival probability estimate (95% C.I.) |  |  |  |
|---|---|---|---|---|
| Group | 2.5 years | 5 years | 7.5 years | 10 years |
| Imipramine | 0.9405 (0.7811–0.9849) | 0.8147 (0.6022–0.9205) | 0.6606 (0.4231–0.8188) | 0.1802 (0.0454–0.3867) |
| Other TCAs | 0.8187 (0.7815–0.8502) | 0.6188 (0.5679–0.6656) | 0.4186 (0.3613–0.4748) | 0.2575 (0.2018–0.3166) |
| Non-TCA | 0.7658 (0.7571–0.7743) | 0.5002 (0.4887–0.5116) | 0.2901 (0.2785–0.3019) | 0.1575 (0.1471–0.1681) |
| No antidx | 0.8130 (0.8052–0.8204) | 0.5907 (0.5796–0.6016) | 0.3898 (0.3773–0.4023) | 0.2477 (0.2352–0.2603) |

A Cox proportional hazards model was also fit to summarize the contrasts between medication groups. The results of the Cox model largely confirmed the results of the KM analysis, with a statistically significant omnibus test, and with imipramine use being associated with reduced hazard of mortality relative to use of a non-TCA, and an estimated hazards reduction relative to no antidepressant use or use of any other TCA (**Table S4**). As described previously, the latter two contrasts are likely underpowered, yet a hazard reduction of 30% would be highly clinically significant. Consistent throughout the analysis was the fact that use of either a TCA or no antidepressants was associated with significantly lower risk of mortality compared to use of a non-TCA antidepressant. Together, these data indicate that NACC participants who used imipramine or a different TCA survived longer, potentially due to a reduced risk for AD-related mortality.

## DISCUSSION

Multiple potential mechanisms by which apoE4 confers increased risk of AD have been proposed and investigated.(49) One hypothesis that we and others have investigated is that apoE promotes the polymerization of Aβ into neurotoxic oligomers and filaments, with apoE4 being the strongest catalyst or pathological chaperone. Substantial evidence from biochemical, transgenic and knockout mouse, and human studies supports this premise and aligns well with the epidemiological evidence.(3-13) First, it was established that apoE binds to the Aβ peptide and promotes its polymerization, and then that apoE expression is essential for amyloid formation in transgenic animal models of AD.(10) Indeed, a report of a human without any *APOE* expression showed no evidence of AD cognitive decline at age 40, which might be expected if apoE were crucial for Aβ clearance from the brain in AD.(50) Thus, the catalytic activity of apoE on Aβ polymerization remains the most likely mechanism by which it confers risk for AD. Interestingly, inflammation and apoE overexpression are features of the AD brain, possibly due to activation of microglia and their secretion of inflammatory cytokines, which stimulate astrocytes to express apoE, thus promoting Aβ polymerization.(10)

In the current study, we found that imipramine use by AD participants in the NACC database led to greatly improved survival over ten years, compared to treatment with other tricyclic, and especially compared to treatment with non-tricyclic antidepressants, and compared to AD participants not prescribed any antidepressant. Some of our results are consistent with findings from a large national AD registry cohort in Sweden, which also found that SSRI/SNRI use was associated with higher risk of all-cause mortality.(51) An essential test of any hypothesis about the mechanism of a human disease such as AD is whether it leads to the finding of a potential intervention that can be tested and shown to be efficacious in human patients. Our previous work and the results reported here indicate that imipramine may satisfy that criterion. A limitation of our study was the small number NACC participants prescribed imipramine, precluding the investigation of sex and race and/or ethnicity as modifying variables, which should be considered in future prospective studies.

Imipramine has been evaluated before in several preclinical AD models where it has been shown to reduce Aβ aggregation and improve cognition.(18, 52-54) We showed here in human AD COs and in transgenic AD rats that imipramine reduces Aβ accumulation and neurodegeneration. The neuroprotective effects of reducing either the concentration or the aggregation state of Aβ in neural tissues are well established, and our current and prior data indicate that both are achieved by imipramine treatment.(18) Besides directly blocking the Aβ-apoE interaction, other indirect effects of imipramine may contribute to its therapeutic benefit, including its effect on glial cell reactivity observed in imipramine-treated human AD COs. Astrocytes uptake and degrade Aβ peptides, processes that have been shown to be potentially mediated by apoE.(55)

That imipramine may improve cognitive deficits via enhanced cholinergic neurotransmission is consistent with prior reports that cholinesterase inhibitors improve spatial working memory deficits in rats.(56) Imipramine can bind directly to AChE and BChE, thereby inhibiting their enzymatic activities.(57) In humans, cholinesterase inhibitors also improve cognition and reduce the risk of AD-related mortality.(58) Despite conflicting mechanisms of action and concerns over its use in the elderly,(19, 59) our data indicate that controlled trials of imipramine combined with a cholinesterase inhibitor to manage anticholinergic side effects may be warranted. Indeed, most AD patients receive cholinesterase inhibitors as standard of care which may obviate the cholinergic side effects of imipramine, revealing its anti-amyloid and pro-survival effects.

Because imipramine targets the essential role of apoE in AD, a clinical trial would be particularly appropriate for participants who carry an *APOE4* allele, who are also less recommended for the current FDA-approved anti-amyloid monoclonal antibody drugs.(14, 15) When taken together with our prior reports, imipramine has now been demonstrated to reduce AD phenotypes in cells, human COs, mice, and rats, and to improve cognition and reduce mortality in human AD patients with depression, with implications for its use as a general treatment for AD.

## CONCLUSIONS

All the available evidence indicates that imipramine use to treat depression in AD patients leads to improved cognition and reduced mortality over ten years compared to other tricyclic or non-tricyclic antidepressants, including SSRIs, arguing for a change in AD clinical practice to consider imipramine as the preferred antidepressant, despite current reluctance to prescribe imipramine to the elderly. Trials to test the effect of imipramine on cognition and mortality in all AD patients, with or without depression, are indicated and are currently being designed.

## LIST OF ABBREVIATIONS

AD: Alzheimer’s disease
SSRI: Selective serotonin reuptake inhibitor
SNRI: Selective norepinephrine reuptake inhibitor CO Cerebral organoid
APOE: Apolipoprotein E
FDA: Food and Drug Administration
AChE: Acetylcholinesterase
BChE: Butyrylcholinesterase
Aβ: Amyloid-β
NACC: National Alzheimer’s Coordinating Center
TCA: tricyclic antidepressant KM Kaplan-Meier
APP: Amyloid precursor protein
HSQC: Heteronuclear single-quantum correlation
FA: Formic acid
MWM: Morris water maze
NMR: Nuclear magnetic resonance
STD: Saturation transfer difference
DPBS: Dulbecco’s phosphate buffered saline
MMSE: Mini-mental State Exam
IHC: Immunohistochemistry
hiPSC: Human induced pluripotent stem cell

## DECLARATIONS

### Ethics approval and consent to participate

All human data from NACC are deidentified. Rats were treated in accordance with the Guide for the Care and Use of Laboratory Animals. All procedures were approved by the Institutional Animal Care and Use Committee of the University of Colorado.

### Consent for publication

Not applicable.

### Availability of data and materials

All human subject data are freely available upon request from NACC. All other datasets supporting the conclusions of this article are included within the article and its additional files.

## Author’s contributions

NRJ and HP were the co-principal investigators. NRJ, RW, LV and HP were responsible for conceptualization and design of the study. NRJ, JB, SS, JR, AW, MMA, DQ, DS, MAH, RW and LV were responsible for data acquisition. NRJ, SS, DS, MAH, RW and LV were responsible for data analysis. NRJ, SS, MAH, BV, HJC, RW, LV and HP were responsible for data interpretation. NRJ, BV, TW, LV and HP supervised the study. NRJ, LV and HP wrote the manuscript. All authors read and approved the final manuscript.

## Acknowledgements

We thank C. Neill Epperson, MD, Chair of Psychiatry, University of Colorado Anschutz, for insightful discussions on the clinical implications of our findings. We thank Julie A. Reisz, PhD, the University of Colorado School of Medicine Metabolomics Core, and the University of Colorado Biophysics Shared Resource Facility for their contributions to the study.

## Clinical trial number

Not applicable.

## SUPPLEMENTARY INFORMATION

Additional file 1. **Figure S1.** One-dimensional trace of NOESY spectrum of imipramine and Aβ42. **Figure S2.** NMR measurements indicate that imipramine does not bind to Aβ40. **Figure S3.** NMR measurements of imipramine binding to Aβ42. **Figure S4.** TgF344-AD rat Morris water maze Day 1 performance metrics. **Figure S5.** Correlation between TgF344-AD Morris water maze (MWM) performance and brain imipramine concentration. **Figure S6.** Immunohistochemical quantification of NeuN staining in the TgF344-AD rat hippocampus. **Table S1.** Rat age and body weight at treatment initiation. **Table S2.** NACC participants in survival and mortality hazards analyses. **Table S3.** Kaplan-Meier survival analyses. **Table S4.** Cox proportional hazard ratio estimates.

Additional file 2. **Data S1. Numbers of participants at risk and censored in Kaplan-Meier analyses.** NACC participants seen at Alzheimer’s Disease Research Centers (ADRCs) who reported being prescribed imipramine, any other tricyclic antidepressant (TCA), any non-TCA antidepressant, or no antidepressants (antidx) were identified. The numbers of participants at risk and censored at each KM time are provided.

