## Additional file 1 for "Imipramine binds to amyloid-β and extends the survival of Alzheimer’s disease patients"

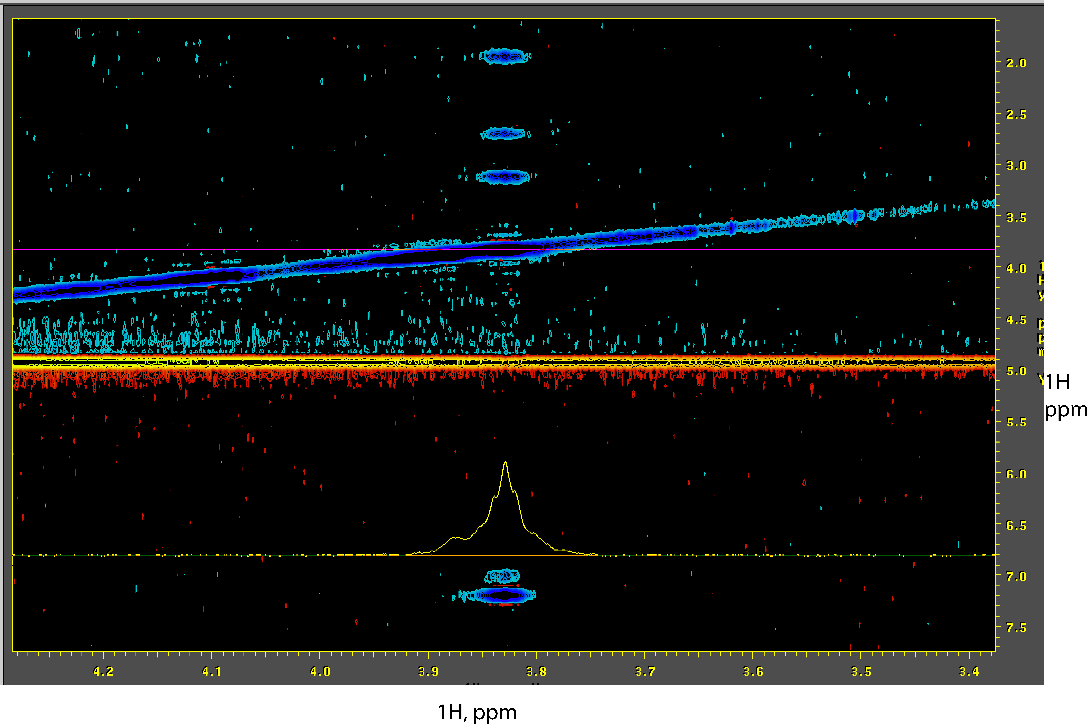

**Figure S1. One-dimensional trace of NOESY spectrum of imipramine and Aβ42.** Zoomed-in region of the ^1^H-^1^H NOESY spectrum of imipramine peaks in the presence of Aβ42. The broadening of the peaks provides additional confirmation of imipramine binding to Aβ42 and increased tumbling time.

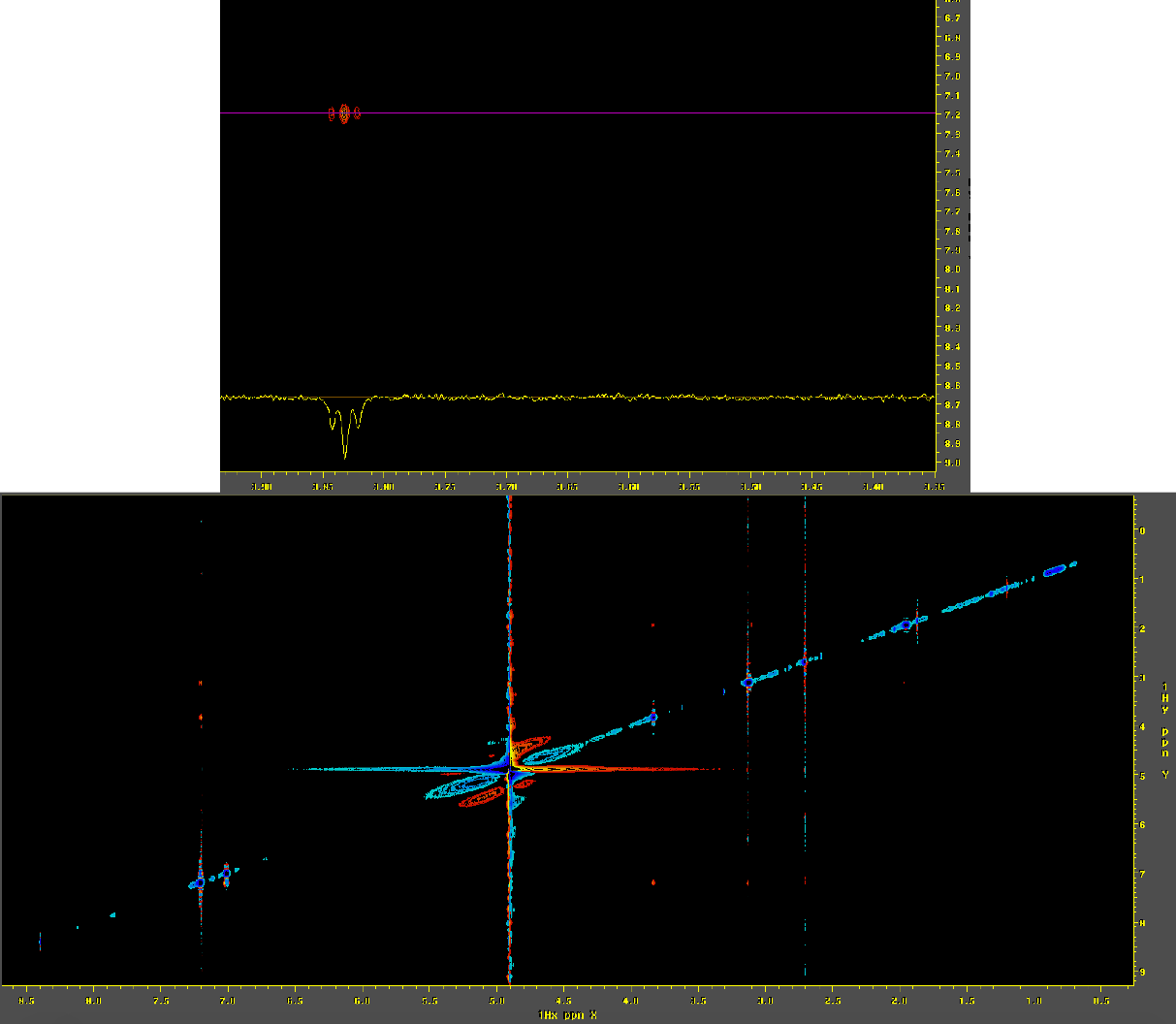

**Figure S2. NMR measurements indicate that imipramine does not bind to Aβ40.** ^1^H-^1^H NOESY spectrum for 200 µM imipramine and 22 µM Aβ40 at 10 °C. The mixing time was 300 ms. The insert on top shows well-resolved imipramine peaks of the opposite sign to the diagonal indicating no binding and a small molecule tumbling behavior.

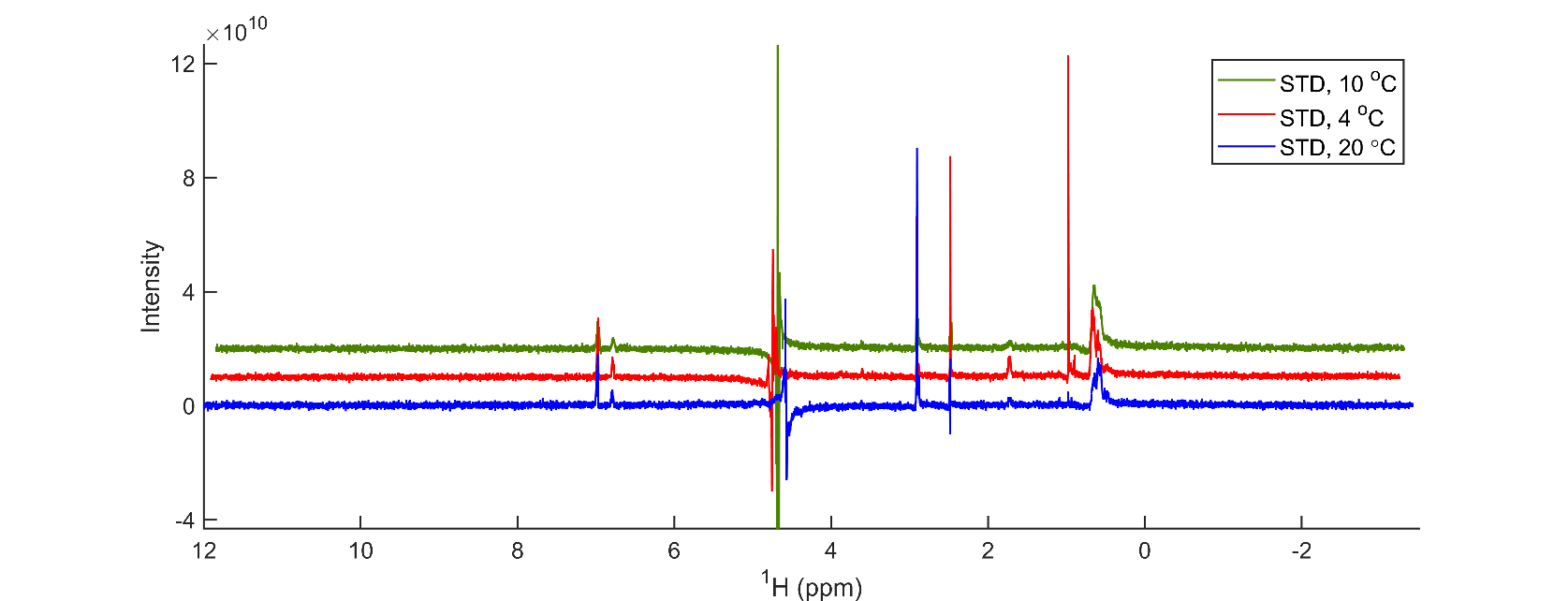

**Figure S3. NMR measurements of imipramine binding to Aβ42.** ^1^H NMR STD spectrum for 200 µM imipramine with 20 µM Aβ42 at 4 (red), 10 (green) and 20 °C (blue).

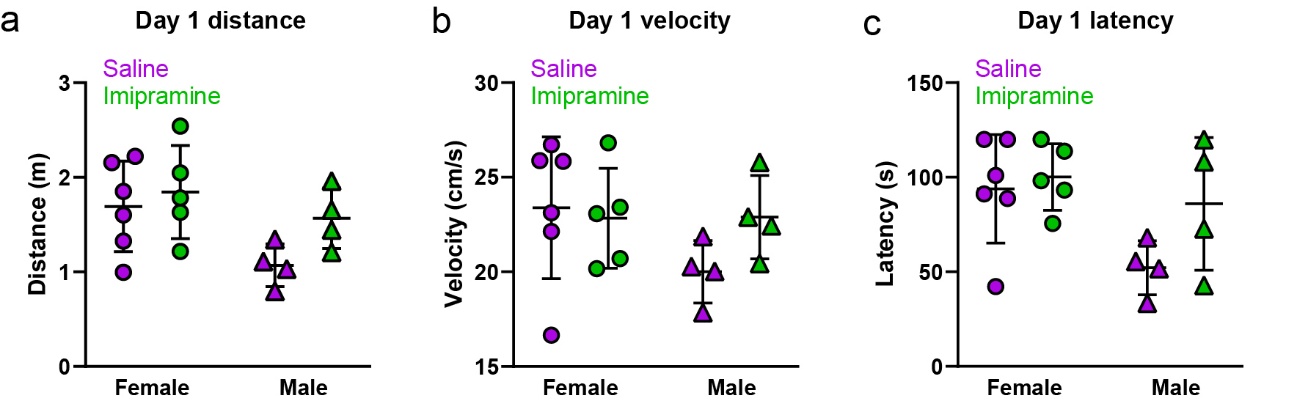

**Figure S4. TgF344-AD rat Morris water maze Day 1 performance metrics.** a) Average distance swum per trial. b) Average swimming velocity. c) Average latency to the platform. Two-way ANOVA.

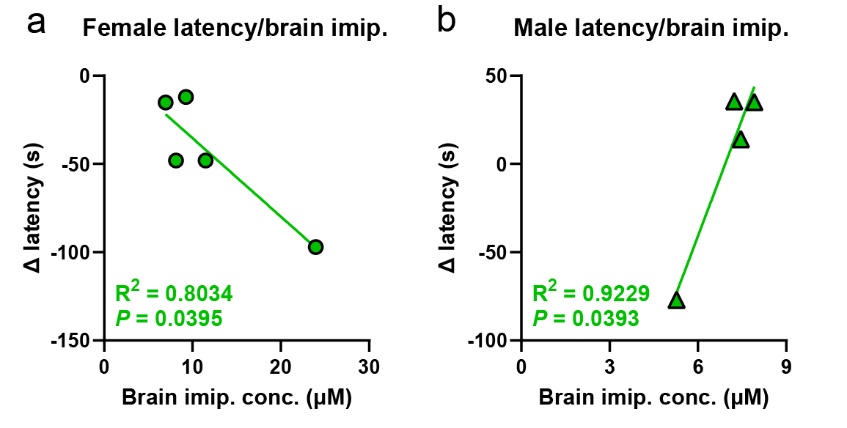

**Figure S5. Correlation between TgF344-AD Morris water maze (MWM) performance and brain imipramine concentration.** a,b) Change in latency to the platform zone between the final training and probe trials versus brain imipramine concentration at sacrifice in imipramine-treated (a) female and (b) male TgF344-AD rats. Simple linear regression and Pearson’s correlation test.

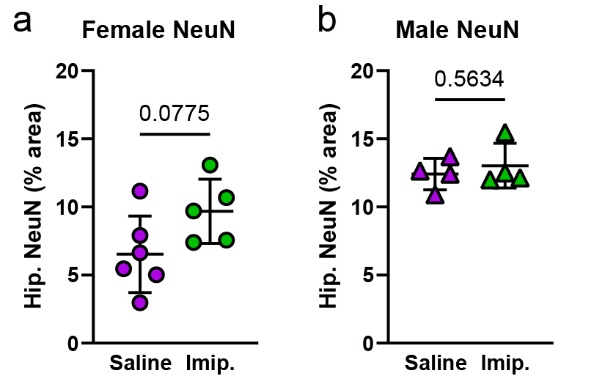

**Figure S6. Immunohistochemical quantification of NeuN staining in the TgF344-AD rat hippocampus.** a,b) Quantification of NeuN^+^ neuronal nuclei area fraction in the hippocampi of saline- and imipramine-treated (a) female and (b) male TgF344-AD rats. Unpaired t-test. Each symbol represents an individual rat. Lines indicate group means ± SD.

| Sex | Treatment | N | Age (mo) | Weight (g) |
| --- | --- | --- | --- | --- |
| Female | Saline | 6 | 16.1 ± 1.5 | 325 ± 73 |
| Female | Imipramine | 5 | 16.0 ± 1.3 | 308 ± 42 |
| Male | Saline | 4 | 20.3 ± 0.5 | 438 ± 34 |
| Male | Imipramine | 4 | 19.8 ± 1.9 | 437 ± 40 |

**Table S1. Rat age and body weight at treatment initiation.** Data indicate mean ± S.D.

| Group | Participants  (#) | Age  (y) Mean ± SD | Female  # (%) | *APOE4**  #/# reported (%) | Deaths  # (%) | Median Exposure Time (y) [IQR] |
| --- | --- | --- | --- | --- | --- | --- |
| Imipramine | 42 | 76.9 ± 10.5 | 25  (59.52%) | 8/35  (22.86%) | 16  (38.10%) | 3.99  [1.48, 7.68] |
| Other TCAs | 596 | 73.1 ± 10.4 | 352 (59.06%) | 206/500 (41.20%) | 240 (40.27%) | 2.93  [1.32, 5.77] |
| Non-TCA | 11,635 | 73.2 ± 10.2 | 6,114 (52.55%) | 4,789/9,875 (48.50%) | 5,705 (49.03%) | 2.66  [1.22, 5.06] |
| No antidx | 13,212 | 75.7 ± 10.1 | 6,690 (50.64%) | 4,645/10,793 (43.04%) | 5,073 (38.40%) | 2.62  [1.11, 5.21] |
| Total | 25,485 | 74.5 ± 10.3 | 13,181 (51.72%) | 9,648/21,203 (45.50%) | 11,034 (43.30%) | 2.64  [1.17, 5.15] |

**Table S2. NACC participants in survival and mortality hazards analyses.** **APOE4* status was not reported for all participants.

| Contrast | Peto-Peto *P* value | Log Rank *P* value |
| --- | --- | --- |
| Overall | <0.0001 | <0.0001 |
| Imipramine vs Other TCA | 0.0450 | 0.2015 |
| Imipramine vs. Non-TCA | 0.0012 | 0.0094 |
| Imipramine vs. No antidx | 0.0263 | 0.1400 |
| Other TCA vs. No antidx | 0.5308 | 0.4716 |
| Other TCA vs. Non-TCA | <0.0001 | <0.0001 |
| Non-TCA vs. No antidx | <0.0001 | <0.0001 |

**Table S3. Kaplan-Meier survival analyses.** NACC participants seen at Alzheimer’s Disease Research Centers (ADRCs) who reported being prescribed imipramine, any other tricyclic antidepressant (TCA), any non-TCA antidepressant, or no antidepressants (antidx) were identified. Kaplan-Meier survival analysis was performed. Exposure time was started when a medication was first recorded, or at their first visit in the case of no antidx, and survival over 10 years was compared by Peto-Peto and Log Rank tests.

| Contrast | Hazard ratio estimate (95% C.I) | *P* value |
| --- | --- | --- |
| Overall | N/A | <0.0001* |
| Imipramine vs. Other TCA | 0.7291 (0.4400–1.2081) | 0.2201 |
| Imipramine vs. Non-TCA | 0.5307 (0.3252–0.8659) | 0.0112* |
| Imipramine vs. No antidx | 0.6949 (0.4258–1.1340) | 0.1452 |
| Other TCA vs. Non-TCA | 0.7279 (0.6397–0.8282) | <0.0001* |
| Other TCA vs. No antidx | 0.9531 (0.8373–1.0848) | 0.4670 |
| Non-TCA vs. No antidx | 1.3094 (1.2608–1.3599) | <0.0001* |

**Table S4. Cox proportional hazard ratio estimates.** Asterisks indicate *P* < 0.05.
